# BC-Predict Database: A Curated Resource of Experimentally Validated Markers in Multidrug Resistance in Breast Cancer

**DOI:** 10.1101/2025.10.09.681460

**Authors:** Sakshi Sanjay Parate, Rahad Rehas, Gupta Soyam, Lia Susan George, Mahammad Nisar, Akshay Unni, TR Sandra, Sophia Manuel, Vineetha Shaji, Aleena Krishna S.V, Mejo George, R Bhadra, Radul R Dev, B Pravin, Siva Subramanian Ayeraselvan, K Majma, TK Ajitha, Ritobrato Chatterjee, VG Rahul, Marlu Jogy, PG Roopashree, Chandhana Prakash, Anagha Muralidharan, Anagha Prakash, Shubham Sukerndeo Upadhyay, Ashna Anilkumar, Niyas Rehman, Manavalan Vijayakumar, Rohan Shetty, Jalaluddin Akbar Kandel Codi, Thottethodi Subrahmanya Keshava Prasad, Anoop Kumar G Velikkakath, Rajesh Raju

## Abstract

**Background:** In this study, we aim to develop a yearly updatable database that could predict chemotherapeutic drug resistance and overall survival probability in breast cancer patients. Existing drug sensitivity databases depend on correlation-based predictions. In our study, candidates involved in drug resistance are chosen based on cell line validation (overexpression or downregulation or inhibition of candidates) studies, curated manually.

**Method:** 28,773 mRNA expression signatures from 914 breast cancer patients were extracted from cProsite. 106 of these patients had clinical information and log2 fold change information required for this study. We categorized these patients into deceased and surviving groups from TCGA. To prepare a database that can predict drug resistance and overall survival, we included mRNAs that were over-expressed in at least 80% of the breast cancer patients and mRNAs over-expressed in deceased and surviving groups. In addition, we also reported breast cancer-associated drug resistance candidates which have been reported in cell-line based studies. The database matrix preparation involved an approximate of 15000 manual searches of cell validated studies. (750 candidates x 20 drugs). The database was validated using a publicly available breast cancer patient proteomics data.

**Results:** Our analysis identified a list of top priority candidates associated with multidrug resistance, categorized based on their resistance to >15 drugs, 5-15 drugs, and 2-4 drugs. Analysis of patient profiles in the database revealed that the number of proteins contributing to drug resistance was high in the poor prognosis category compared to the good prognosis category.

**Conclusions:** Our study highlights the probable gaps in breast cancer drug resistance research, as only a small subset of overexpressed mRNA candidates found in patients are studied in vitro or in vivo experiments focusing on drug resistance. We also identified candidates involved in multidrug resistance, whose role in drug resistance has not been studied in more than 15 drugs. After further validations, this will benefit the clinicians and upcoming CRISPR gene therapeutics.

## Introduction

According to Globocan 2022, the incidence and mortality rates of female breast cancer remain high in developed as well as developing countries, including India (1). While genetics contribute to 5-10% as a risk factor for breast cancer (2), other factors such as age, lifestyle, early age of menstruation, late menopause, and obesity also increase the susceptibility (3,4). The World Health Organization established the Global Breast Cancer Initiative (GBCI) in 2021 to overcome the challenges of breast cancer, including early and timely diagnosis, along with therapeutic management (5).

The treatment modalities of breast cancer have progressed over the years, but further research is required for personalized therapy. The typical prognosis for breast cancer is decided based on the Tumor, Node, and Metastasis (TNM) breast cancer staging system: Stage 0, Stage I, Stage II, Stage III, and Stage IV, histology, and biomarkers. Present treatment strategies include tumor surgery, radiation, targeted therapy, immunotherapy, and endocrine therapy to focus on the breast conservation approach or mastectomy (5,6). Stages I and II breast cancers are usually treated with breast-conserving surgery and radiation therapy. Radiation therapy following breast-conserving surgery decreases mortality and recurrence (7). Chemotherapy remains the standard of care for the majority of invasive and metastatic breast cancer treatment, but unfortunately, patients frequently develop resistance. This poses a major challenge in the management of breast cancer.

Over time, the tumor cells try to evade the effects of chemotherapy agents, leading to treatment failure and disease progression. The drug resistance can emerge through different mechanisms including the involvement of cancer stem cells (CSCs), genetic and epigenetic factors, tumor microenvironment, drug transport, target binding efficiency, cancer-associated fibroblasts (CAFs), drug efflux, upregulation of multi-drug resistance proteins, altered drug targets, apoptotic inhibition, aggressive EMT characteristics, pH alteration, altered cellular metabolism, autophagy, increased DNA repair, and involvement of miRNAs (8,9).

Several databases and tools provide information on the lifespan of cancer patients based on genomics or proteomics data. Some of the prominent databases include The Cancer Genome Atlas (TCGA), cBioPortal for Cancer Genomics, Gene Expression Omnibus Database (GEO), PROGgeneV2, Clinical Proteomic Tumor Analysis Consortium (CPTAC), University of

ALabama at Birmingham CANcer (UALCAN), Human Cancer Proteome Variation Database (CanProVar), and drug resistance databases such as Cancer Drug Resistance Database (CancerDR) (10), DRMref (11), Drug resistance information (DRESIS) (11,12), and HNCDrugResDb (13). These databases and tools offer valuable resources for researchers studying the relationship between genomics, proteomics, and cancer prognosis. However, it’s important to note that predicting lifespan in cancer patients is complex and influenced by various factors beyond genomic and proteomic profiles, such as treatment response, lifestyle factors, and comorbidities (14).

Genomic and transcriptomic approaches have proven instrumental in developing potential biomarker panels for breast cancer. Techniques such as whole exome sequencing (WES) provide information about the tumor mutational burden (TMB). A lot of advanced combination endocrine therapy has proved beneficial for breast cancer patients. However, some of them develop resistance, and the markers to predict the resistance are still lacking (15). This is majorly due to the inter- and intra-heterogeneity of breast cancer (15)(16). As multiple population studies are required to arrive at better conclusions and improve drug resistance predictions, here we contribute several additional pieces of information to this field.

In this study, we identified overexpressed mRNA profiles in tumor tissues compared to adjacent normal tissues of breast cancer patients (extracted from cProsite). By analyzing 106 breast cancer patients, we identified a subset of genes overexpressed in at least 80% of this cohort. We also identified mRNA overexpression signatures that distinguish deceased vs surviving groups. These were further manually curated against a panel of 20 anticancer drugs (17)(18) to identify their role in drug resistance. To expand our dataset, we integrated the molecules responsible for drug resistance from the published breast cancer cell line-based studies. We also identified candidates that can cause multidrug resistance against multiple chemotherapy drugs if differentially expressed. We compiled all the information to build a database called BC-Predict Database. Drug resistance patterns and survival probability predicted for the breast cancer patients based on their mRNA profiles are demonstrated.

## Methodology

### Extracting mRNA expression data of breast cancer patients from cProSite

We extracted expression data for 28,773 mRNAs from 914 breast cancer patients using Proteogenomic Data Analysis Site (cProSite) website (19), a public cancer resource developed by the National Cancer Institute (NCI). Among these patients, 106 patients fulfilled the criteria for further analysis as they had sufficient clinical information in addition to tumor abundance and adjacent normal tissue expression values as log2 fold change (log2FC). For extracting the data, we applied ‘Breast Cancer’ as the Tumor type in the cProSite interface, and ‘RNA level’ as the Dataset and selected ‘Tumor vs Normal Tissue’ for Analysis. The Cancer Genome Atlas (TCGA) (https://www.cancer.gov/tcga) was then chosen from the Tumor View module for each gene, culminating into a single file for further analysis.

### Categorizing breast cancer patients based on their survival data from the TCGA database

The breast cancer patient list obtained from cProsite was cross-checked with The Cancer Genome Atlas (TCGA) database. The patient’s clinical information, including survival data, was extracted. Based on the TCGA data, the 106 patients were segregated into two main groups: deceased and surviving. This was done using a custom script to automate the data retrieval process through cProsite’s application programming interface (API). The mRNA expression data was compiled into a tab-delimited (TSV) file to carry out further analysis. Further, IBM SPSS Statistics Version 27 was used to determine statistical significance from mRNA expression profiles of deceased vs surviving patient groups. Nonparametric tests were conducted for independent samples in a customized manner using log2FC values. Mann-Whitney U Test of independent samples was used to test null and alternative hypotheses. We then filtered and sorted the data based on the significant p-value of ≤0.01 and corresponding log2FC. The top 50 upregulated mRNAs from both deceased and surviving patients were further studied for their involvement in drug resistance using manual curation.

### Inventory of differentially expressed mRNA candidates present in the majority of breast cancer patients

The log2FC values obtained from cProsite was used to identify the mRNAs differentially expressed in at least 80% of the 106 breast cancer patients. The mRNAs with a log2FC of greater than 1.5 were considered significantly upregulated or overexpressed, and those less than -1.5 were considered significantly downregulated. Heatmaps were generated to visualize the expression patterns of the differentially expressed mRNAs across the patient cohort. The upregulated mRNAs observed in this study were taken as candidates for the drug resistance curation.

### Manual biocuration of molecular alterations associated with breast cancer resistance

The list of candidate mRNAs upregulated in at least 80% breast cancer patients’ population was identified to check for their roles in drug resistance, if any. Furthermore, the list of mRNAs that distinguish the deceased and surviving groups was also included. Along with these candidate mRNAs, other molecules involved in breast cancer drug resistance were also selected based on extensive literature review. A PubMed search was conducted to retrieve research articles on breast cancer drug resistance. This retrieved a list of molecules which was further divided into three breast cancer subtypes: Luminal (MCF7- ER+, PR+, HER2), HER2 positive (MDAMB453- ER-, PR-, HER2+), and Triple-negative breast cancer (TNBC) (MDAMB468- ER-, PR-, HER2-). Cell lines specific to these subtypes were selected. The search term used for the biocuration of chemotherapy drug resistance molecules included: “(MCF7 OR MCF-7)” AND “drug resistance” NOT “review” for luminal subtype, “(MDAMB453 OR MDA-MB-453)” AND “drug resistance” NOT “review” for HER2 positive subtype, and “(MDAMB231 OR MDA-MB-231)” AND “drug resistance” NOT “review” for TNBC subtype. The molecules included proteins, miRNAs, and lncRNAs. Review articles, editorials, case studies and patient studies were excluded from the curation. The statement from the articles claiming the overexpression or downregulation of a candidate induced drug resistance were included for biocuration to prepare the database. The claim lines were pasted in a spreadsheet along with their corresponding PMIDs, for each protein/candidate along with the names of the drugs to which it induces resistance (**Supplementary Table 6**). Drug names were manually searched for 20 breast cancer drugs available in the market including Doxorubicin (Dox), Cisplatin (Cis),

Cyclophosphamide (Cyc), Carboplatin (Cis), Paclitaxel (Pac), Imatinib (Ima), Docetaxel (Doc), Trastuzumab (Tra), Methotrexate (Met), Rituximab (Rit), Gemcitabine (Gem), Bortezomib (Bor), Vincristine (Vin), Dexamethasone (Dex), Gefitinib (Gef), Tamoxifen (Tam), 5-Fluorouracil (5-FU), Etoposide (Eto), Capecitabine (Cap), And Cytarabine (Cyt). In conditions where the breast cancer studies were not available for a candidate to show its role in drug resistance, we considered studies done in other cancer cell line models. These biocuration results were used for database generation. The search term used for searching the molecule-specific drug resistance included “molecule name” AND “drug name resistance” NOT “review”.

### Identification of candidates involved in multidrug resistance

For the identification of candidates involved in multidrug resistance, we represented the biocuration in the form of a heatmap. The candidates were categorized based on their resistance profiles. This classification of multidrug-resistant candidates consisted of molecules resistant to >15 drugs, 5-15 drugs, and 2-4 drugs.

### Preparation of the breast cancer drug resistance database BC-Predict Database

**BC-Predict Database** is an online web application developed using the Django REST Framework and React.js. The application consists of back-end and front-end components. The back-end of **BC-Predict Database** is a Django application, a powerful and flexible toolkit built with Python for developing web APIs. The front-end is developed using the React JavaScript library, and data visualization is implemented using D3.js. The application is containerized using Docker and deployed on AWS, ensuring scalability, reliability, and ease of maintenance. The database will recognize input files of patients that contain differentially expressed candidates (mRNAs, proteins, miRNAs, and lncRNAs). Candidates involved in drug resistance will be detected, and a pattern will be generated to give an overall picture of multidrug resistance probability.

## Results

### Assembly of a drug resistance database matrix by integration of patient molecular signatures and literature-curated molecules

Based on the chemotherapeutic drugs available in the market, we selected 20 drugs for the biocuration of molecules implicated in breast cancer drug resistance for database assembly. The candidates belonging to 3 different categories were selected for the manual biocuration. These candidates included (1) mRNAs overexpressed in at least 80% of the breast cancer patients obtained from cProsite, (2) top 50 mRNAs overexpressed in the deceased and surviving groups respectively, and (3) candidates including proteins, miRNAs, and lncRNAs already associated with breast cancer drug resistance (resistant against atleast one drug) based on published literature (belonging to luminal, TNBC, and HER2 positive subtypes of breast cancer) were chosen (**Figure 1**). Manual biocuration (involving 15,080 manual searches) was performed for these candidates to identify their roles in drug resistance against 20 chemotherapeutic drugs available in the market such as Doxorubicin (Dox), Cisplatin (Cis), Cyclophosphamide (Cyc), Carboplatin (Cis), Paclitaxel (Pac), Imatinib (Ima), Docetaxel (Doc), Trastuzumab (Tra), Methotrexate (Met), Rituximab (Rit), Gemcitabine (Gem), Bortezomib (Bor), Vincristine (Vin), Dexamethasone (Dex), Gefitinib (Gef), Tamoxifen (Tam), 5-Fluorouracil (5-FU), Etoposide (Eto), Capecitabine (Cap), and Cytarabine (Cyt). The classification of these drugs and their targets is depicted in **Supplementary** Figure 1 **and Supplementary Table 1**.

**Figure 1:**
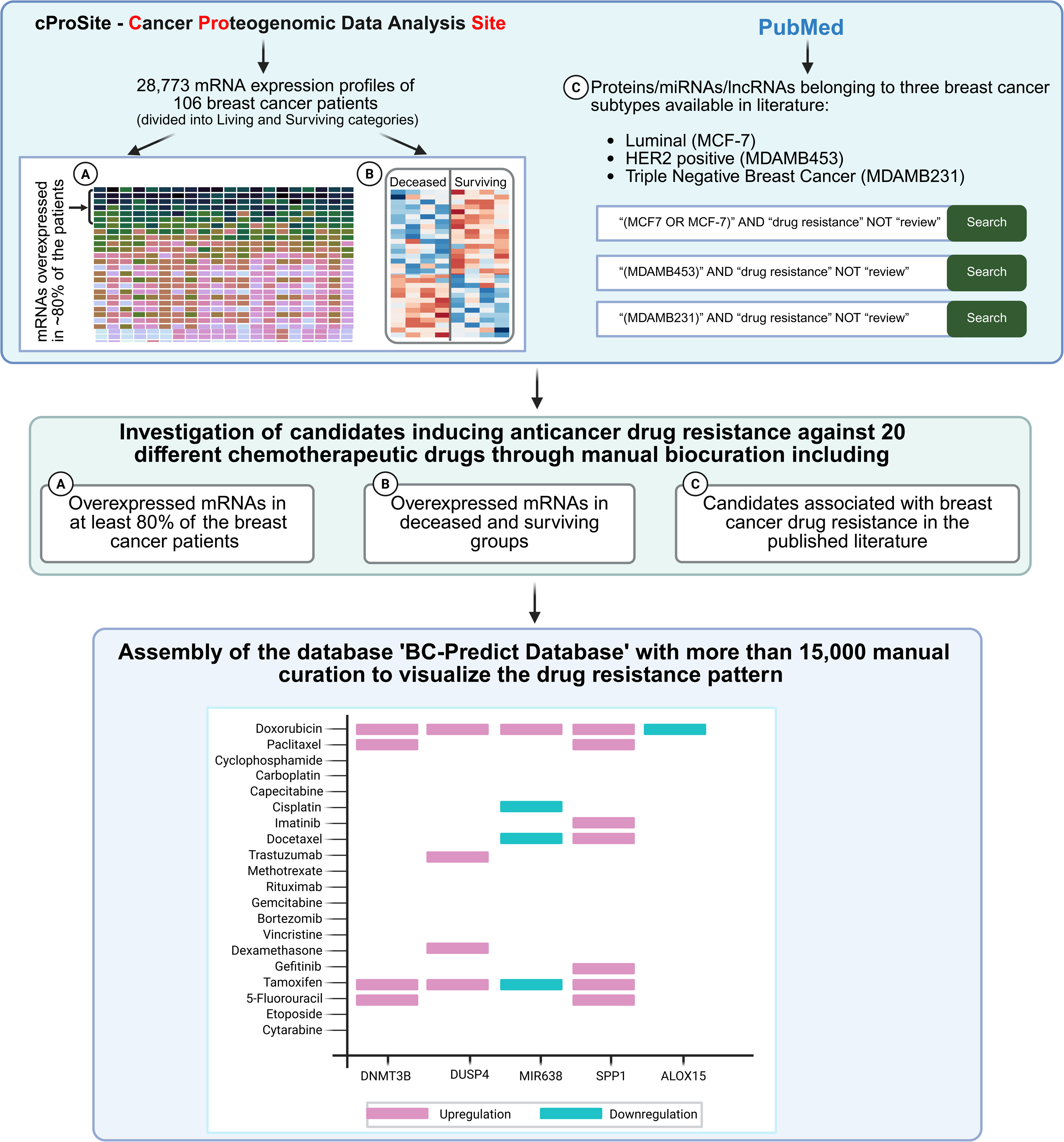
**Schematic presentation of the overall framework employed for the assembly of the drug resistance database**

### Framework for the extraction of breast cancer patient mRNA signature

To assemble a database for visualizing anticancer drug resistance, we extracted mRNA expression profiles of 106 breast cancer patients from cProSite. We obtained 28,773 mRNA expression signatures of these patients. From this cohort of 106 breast cancer patients, 44 belonged to the deceased group and 62 belonged to the surviving group. Additional information on the breast cancer patients extracted from TCGA data is given in **Supplementary Table 2**.

#### 1. mRNAs overexpressed in at least 80 percent of the breast cancer patients

From 28,773 mRNAs obtained from cProsite, 467 mRNAs were identified as significantly overexpressed candidates after applying a log2FC of ±1.5 and they were commonly observed in at least 80% of the patients. Among these, 174 mRNAs were found to be significantly upregulated, and 293 mRNAs were downregulated. The expression profiles of these candidate mRNAs are represented as a heatmap in **Figure 2**. This allows for gaining insights into the gene expression levels across breast cancer patients. The majority of the upregulated mRNAs were involved in cell cycle signaling including mitosis and meiosis, microtubule assembly and reorganization, mitotic spindle organization, chromosome organization, cellular processes regulation, and DNA repair such as CDC6, AURKA, MYBL2, BIRC5, BUB1, CCNE2, CDK1, NUSAP1, NEIL3, FOXM1, MMP13, to name a few represented in **Supplementary** Figure 2 and the list is provided in **Supplementary Table 3**. Some of the mRNAs were also found to be downregulated in at least 80% of the patients. The majority of them were involved in receptor tyrosine kinase signaling, muscle contraction, cardiac contraction, and retinoid metabolism that included FOSB, GFAP, EGFR, VEGFD, FGF17, NRG1, ALDH1L1, SCN2B, ATP1A2, MYH11, and so on (**Supplementary** Figure 3**; Supplementary Table 4**).

**Figure 2.**
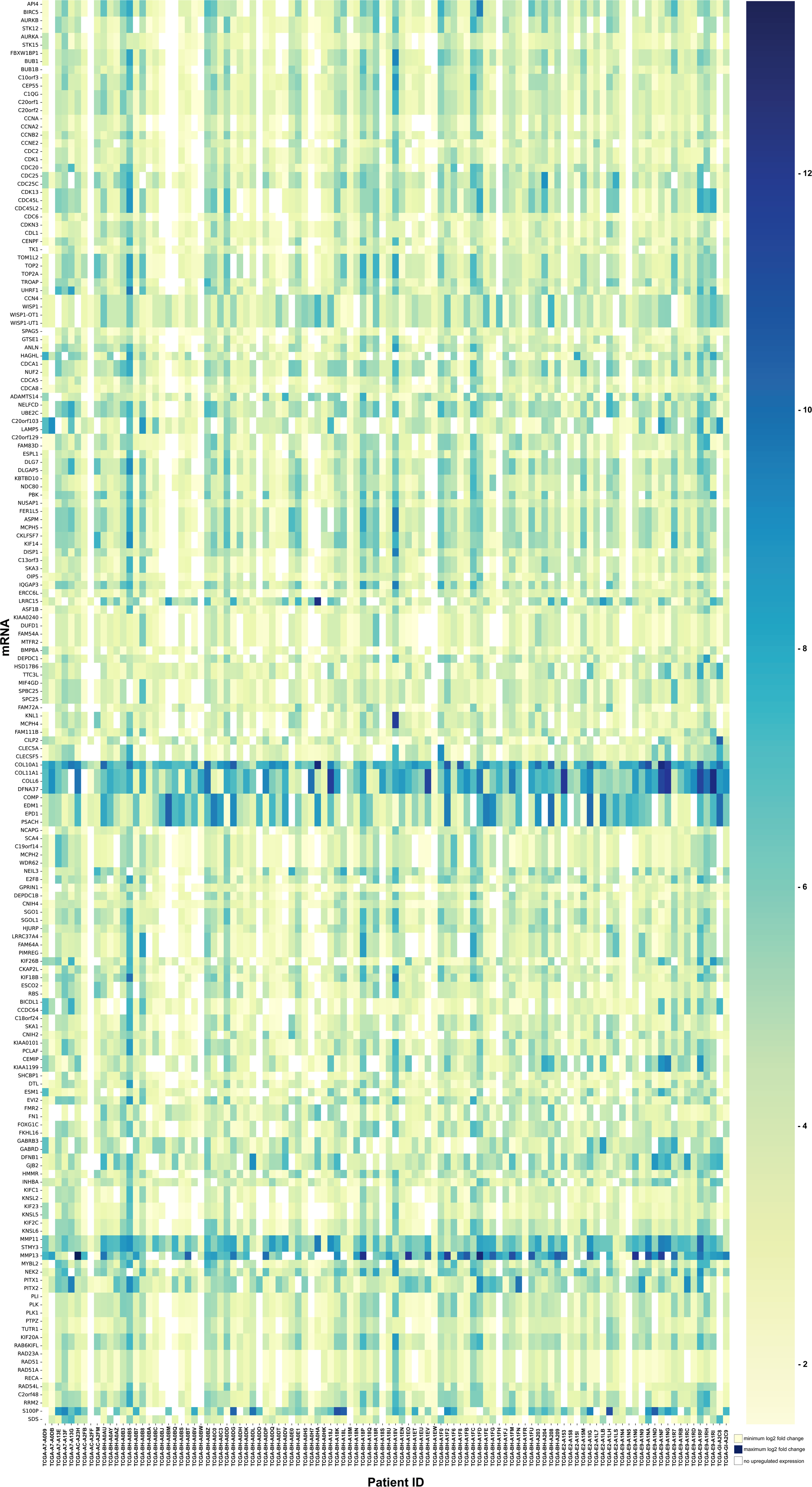
**Heatmap representing the expression patterns of 174 overexpressed candidate mRNAs in atleast 80% of the patients**. Out of the 28,773 mRNAs, 174 were found to be significantly overexpressed in at least 80% of the breast cancer patients. The selection of the candidate mRNAs was done using a log2 fold change cutoff of ±1.5 obtained from the mRNA expression data between the tumor abundance and normal adjacent tissue from cProSite. Each row represents a specific mRNA, and each column depicts a patient sample.

#### 2. Top mRNAs overexpressed in the deceased and surviving groups

The mRNA expression profiles of the breast cancer patients extracted as mentioned above were used here with an aim of identifying mRNA signatures that distinguish deceased vs surviving groups of breast cancer patients. The life expectancy data of these patients were acquired from the TCGA database. They were grouped into two categories based on their survival: 44 deceased and 62 patients in surviving groups. A total of 895 out of 28,773 mRNAs were found to be statistically significant among the breast cancer patients from cProSite. From the SPSS analysis, these mRNAs represented a p-value of <0.01, and rejected the null hypothesis. Their expression signatures were represented as log2FC values between tumors and adjacent normal tissues (**Supplementary Table 5**). A detailed understanding of the significant mRNA expression patterns of deceased and surviving breast cancer patients obtained from cProSite is represented as a combination of the heatmap and bar plots. The heatmap clearly showed a trend of contrasting expression patterns, which confirmed that most of the mRNAs with higher log2FC in deceased and lower log2FC in surviving groups and vice versa, as represented in **Figure 3**. This was complemented with bar plots representing the expression of each gene in the deceased and surviving categories to provide a more focused examination of individual mRNAs (**Supplementary Table 6 & 7)**. The expression of 468 mRNAs out of 28,773 were found to be higher in deceased patients compared to the surviving patients. Top 32 mRNAs from these are represented as bar plots in **Figure 4**. While 417 mRNAs had higher expression in surviving patients compared to deceased patients, the top 32 mRNAs from this group are represented in **Figure 5**. This combined approach provided both global visualizations as well as detailed view of individual gene levels between the two patient categories. To investigate their potential role in drug resistance, we manually curated the top mRNAs in each category. This allowed us to investigate the probable role of these signature mRNAs in drug resistance, if any.

**Figure 3.**
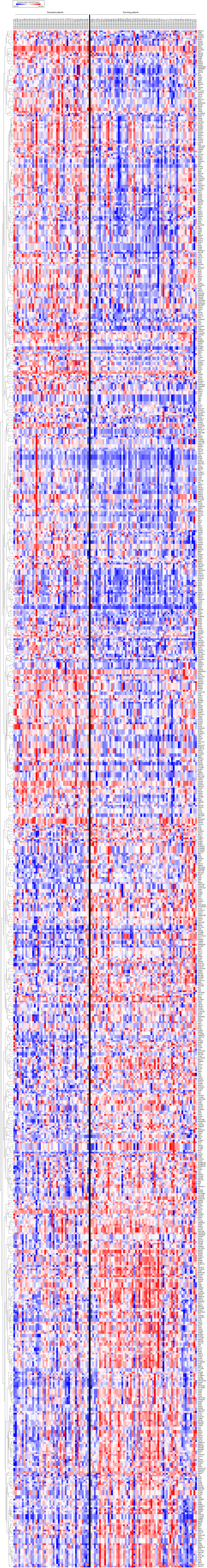
Heat map distinguishing the significant and differentially expressed mRNAs in deceased vs surviving breast cancer patients. The expression of 895 mRNAs is represented in a color gradient (low to high) clustered into two groups: 44 deceased breast cancer patients (left) vs 62 surviving breast cancer patients (right). TCGA patient IDs extracted from cProSite are represented on the top and each column indicates mRNA levels of a single patient. TCGA-A7-A13E to TCGA-E9-A1RB is the deceased category, and TCGA-A7-A0D9 to TCGA-GI-A2C9 is the surviving category. The rows indicate mRNAs involved in deceased vs surviving patient categories. Hierarchical clustering was performed to group the mRNAs depicting similar expression profiles with the resulting dendogram represented on the left.

**Figure 4.**
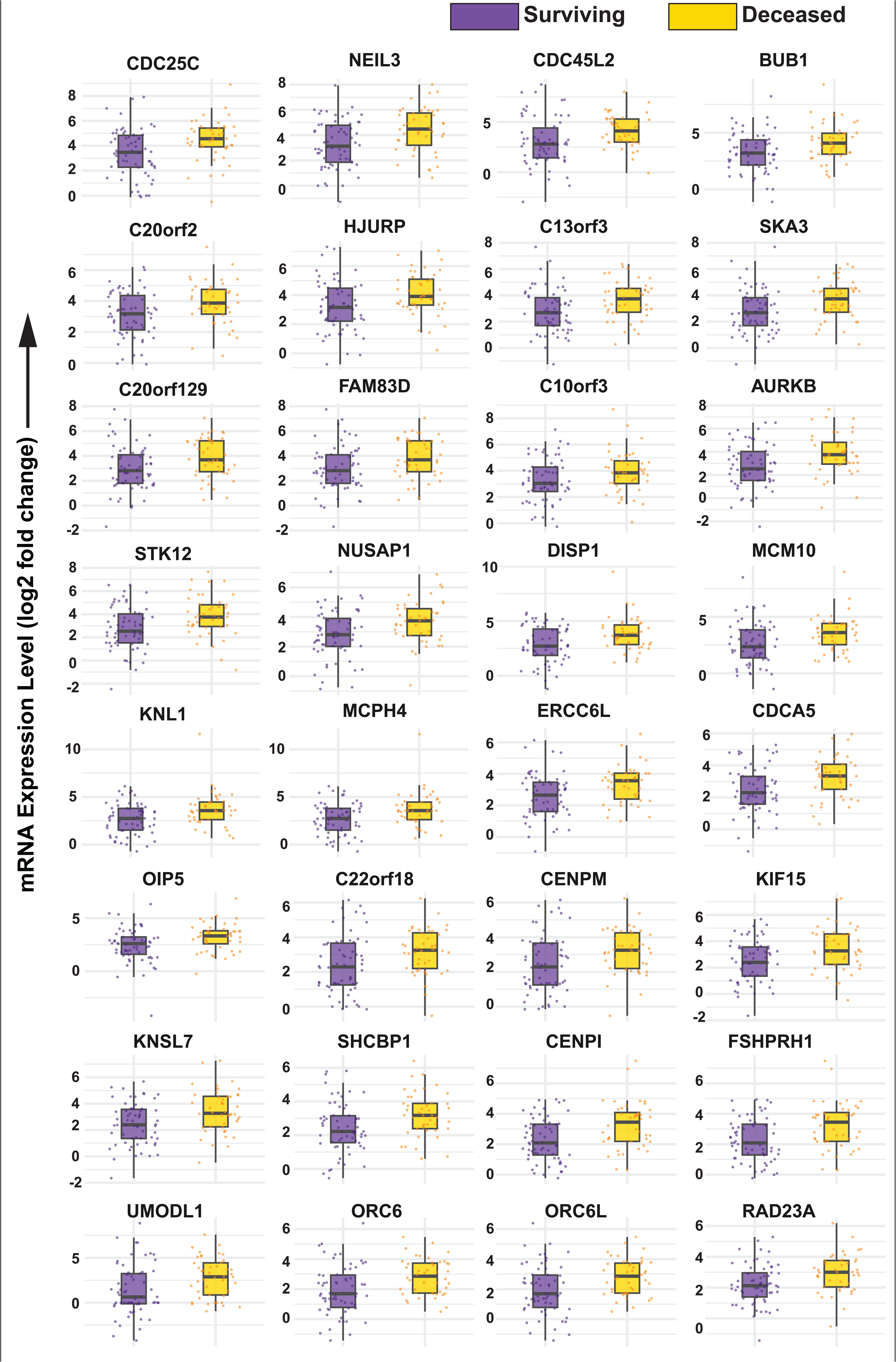
Expression of top 32 mRNAs with higher expression in the deceased category compared to the surviving category. The mRNA expression profiles of each gene are plotted as log2 fold change values calculated between the tumor abundance and adjacent normal tissue obtained from cProSite. It was plotted and represented as bar plot to observe the difference in the deceased versus surviving cohort categories. Top 32 (out of 468) mRNAs are represented with higher expression in the deceased (yellow color) category compared to the surviving (purple color) category.

**Figure 5.**
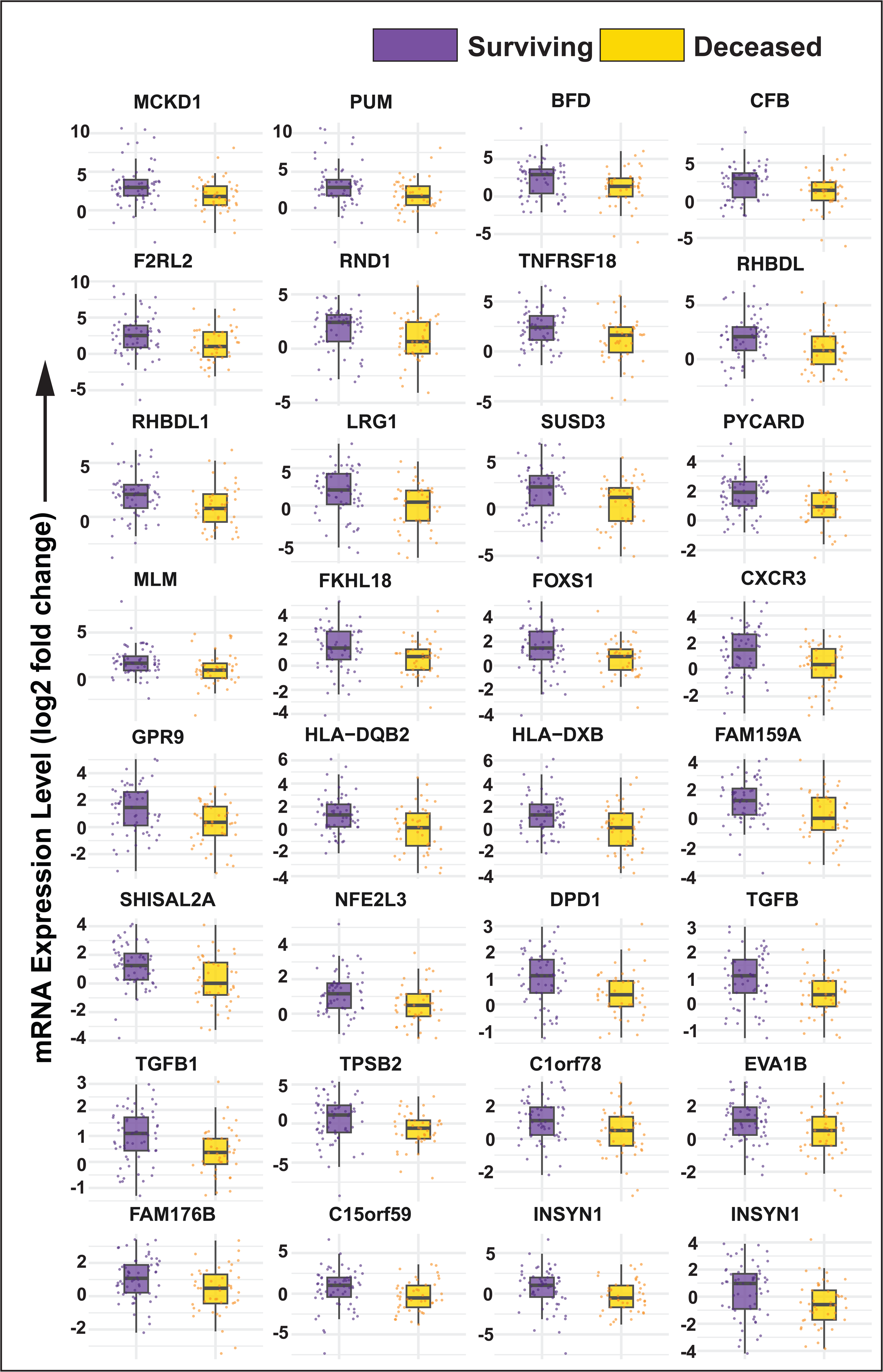
Expression of top 32 mRNAs with higher expression in the surviving category compared to the deceased category. The mRNA expression profiles of each gene are plotted as log2 fold change values calculated between the tumor abundance and adjacent normal tissue obtained from cProSite. It was plotted and represented as bar plot to observe the difference in the deceased versus surviving cohort categories. Top 32 (out of 417) mRNAs are represented with higher expression in the deceased (yellow color) category compared to the surviving (purple color) category.

#### 3. Candidates already associated with breast cancer drug resistance based on published literature

Along with these mRNA candidates, other candidates, including proteins, miRNAs, and lncRNAs previously reported to be involved in resistance against at least one anticancer drug (breast cancer cell line-based studies), were included. These candidates were biocurated against 20 anticancer drugs. Three representative cell lines; MCF-7, MDAMB453, and MDAMB231, corresponding to the three subtypes of breast cancer; luminal, HER2-positive, and triple-negative breast cancer (TNBC), respectively, were selected for this biocuration.

### Identification of molecules known to play a role in drug resistance

Preparation of matrix for the drug resistance database: 750 non-redundant candidates from the above-mentioned categories were curated for their drug resistance pattern against 20 chemotherapy drugs (**Supplementary Table 8**). These 15,000 manual searches (750 candidates x 20 drugs) built the basic matrix for the development of a drug resistance prediction database (**Figure 6a)**. We further examined this matrix and divided it into three groups including 346 candidates showing exclusive upregulation or overexpression scored as 1 (**Figure 6b**), 77 candidates with exclusive downregulation scored as 2 (**Figure 6c**), and 131 candidates showing both overexpression and downregulation for different drugs (**Figure 6c**). The candidates that are tested for their role in drug resistance (by overexpression or silencing or targeted inhibition in cell lines), formed the basis for this database. A mere change in gene expression after addition of a drug was not considered here, unless they were validated by the mentioned criteria for their roles in drug resistance (**Table 1**).

**Figure 6.**
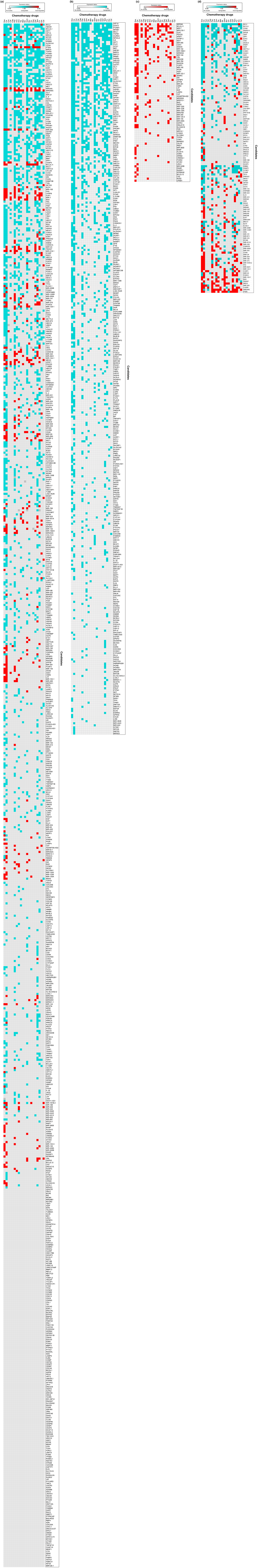
Representation of the biocuration matrix for the anticancer drug resistance database. This matrix represents the biocuration of the overexpressed mRNAs in at least 80% of patients, the mRNAs upregulated in deceased and surviving cohort patients, along with other molecules such as proteins, miRNAs, and lncRNAs involved in breast cancer drug resistance. The x-axis represents 20 chemotherapy drugs and the y-axis represents the targets resistant to these drugs. The light grey color box indicates no studies found for that candidate against that drug. In contrast, the blue color box indicates upregulation of that molecule causing drug resistance and the red color box indicates downregulation of the molecule causing drug resistance. The matrix is divided into four sections: Overall representation of total of 750 candidates in all the categories, **(b)** 346 candidates exclusively upregulated and contributing to drug resistance, **(c)** 77 candidates exclusively showing downregulation contributing to drug resistance, **(d)** 131 candidates showing both upregulation and downregulation for different drugs contributing to drug resistance.

**Table 1:**
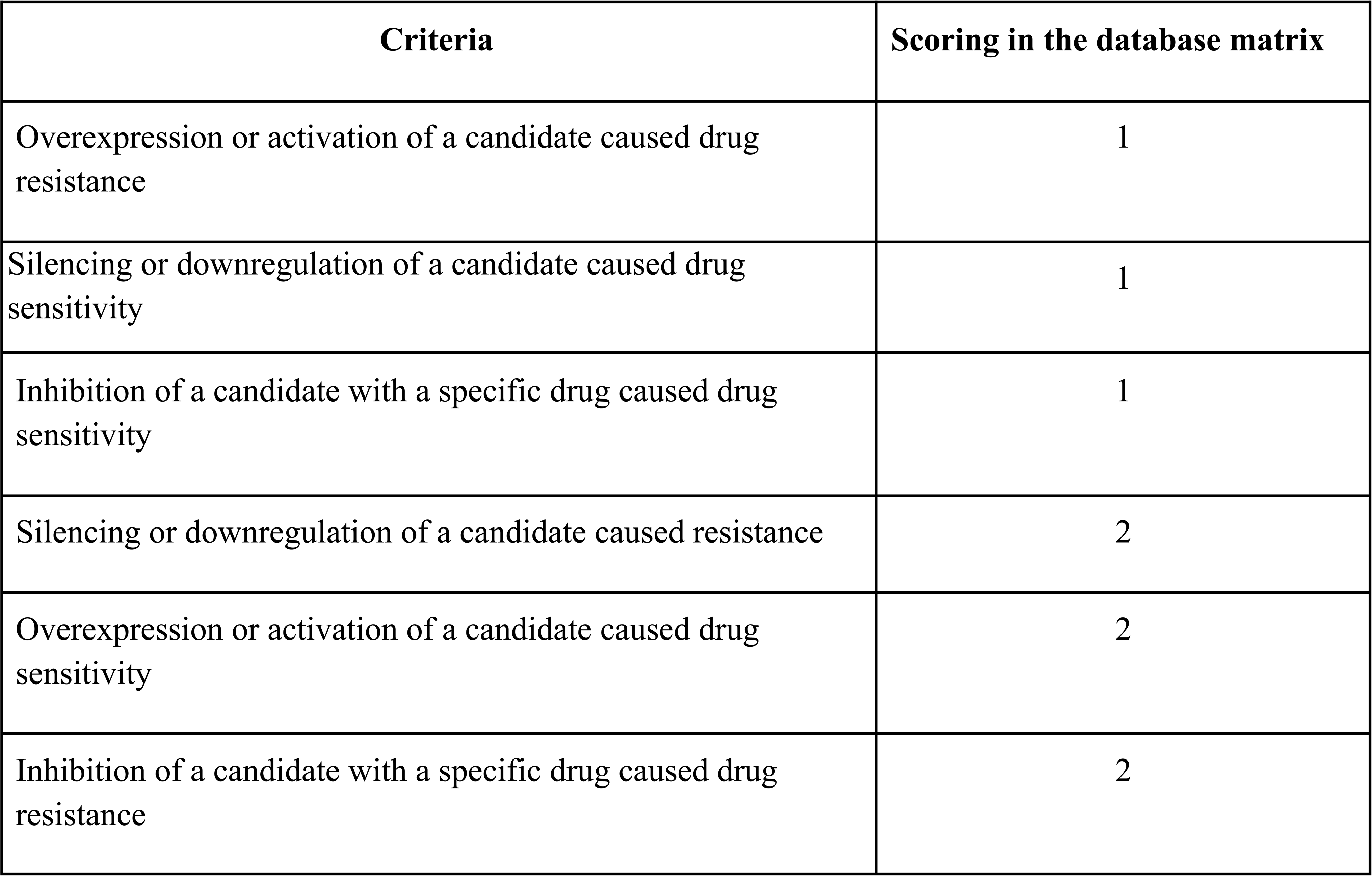
Scoring criteria used for preparation of database matrix.

### Candidates representing multi-drug resistance patterns

After the compilation of drug-resistant candidates, a list of these molecules showing multi-drug resistance was made. Our dataset consisted of a total of 750 molecules from three lists: mRNAs upregulated in at least 80% of patients, mRNAs overexpressed in deceased and surviving groups and molecules reported to be involved in resistance against atleast one drug, in three breast cancer subtypes. We found 566 out of 750 candidates in conferring drug resistance. This included 348 candidates showing resistance to Cis, likewise, 320 to Dox, 239 to Pac, 207 to Tam, 176 to Gem, 165 to FU, 144 to Doc, 106 to Gef, 93 to Eto, 92 to Tra, 75 to Ima, 72 to Bor, 71 to Vin, 70 to Cyt, 66 to Car, 48 to Met, 34 to Dex, 27 to Cyc, 11 to Rit, and 6 to Cap (**Figure 7a**).

**Figure 7:**
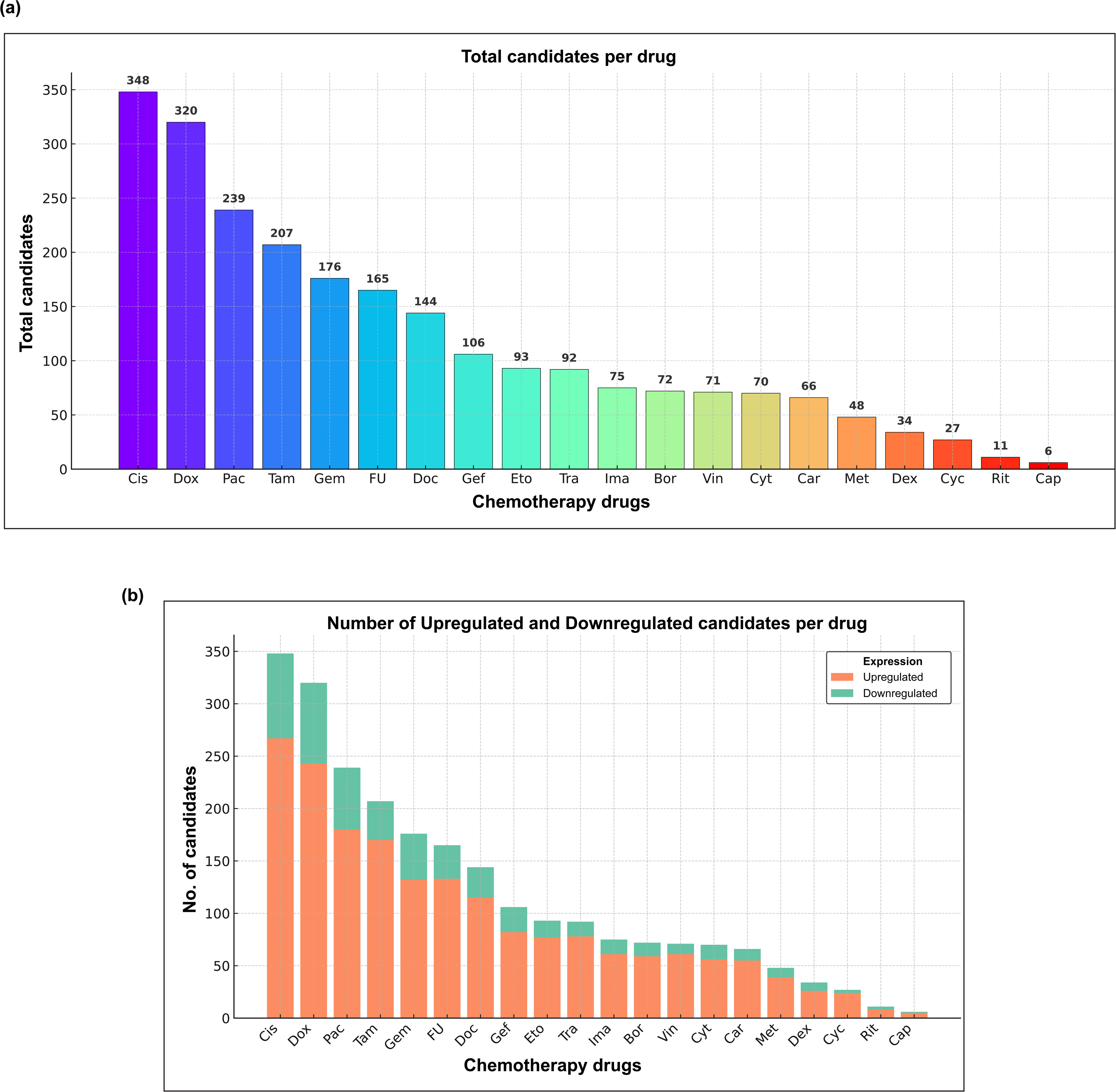
**Number of altered candidate molecules associated with resistance to each of the 20 chemotherapy drugs in the database.** (a) Overall representation of the number of candidates as bar plot involved in resistance to that particular drug, (b) Number of candidates upregulated and downregulated associated with the resistance to that particular drug where the x-axis represents anticancer drugs and y-axis denotes the number of candidates.

This data can be further divided based on the number of candidates upregulated and downregulated, resistant to the particular drug represented in **Figure 7b** (**Supplementary Table 9**).The molecules showing multi-drug resistance were divided into three groups based on the number of resistant drugs. Among these, 11 molecules were involved in resistance to >15 drugs, similarly, 195 molecules were involved in resistance to 5-15 drugs, and 251 were involved in resistance of 2-4 drugs. This data was used to create a framework for visualizing the drug resistance database (**Supplementary Table 10**). The candidates with a potential of causing multidrug resistance against >15 chemotherapy drugs mainly involved in apoptosis inhibition (eg. BCL2), DNA repair (eg. BRCA1), hypoxia signaling (eg. HIF1A). The ABC transporters that play a major role in drug efflux such as ABCG2 was observed to be resistant to 10 drugs out of 20 including Cyc (20), Pac (21), Ima (22), Doc (23), Tra (24), Met (25), Gem (26), Gef (27), FU (28), and Eto (29). Certain proteins such as BCL2 that play a role in apoptosis inhibition and contribute to therapy resistance was found to be resistant to 17 drugs including Dox (30), Cis (31), Cyc (32), Car (33), Pac (34), Ima (35), Doc (36), Rit (37), Gem (38), Bor (39), Vin (40), Dex (41), Gef (42), Tam (43), FU (44), Eto (45), and Cyt (46). Some EMT markers, such as ZEB1, showed resistance to Dox (47), Cis (48), Pac (49), Doc (50), Tra (51), Met (52), Gem (53), Gef (54), Tam (55), FU (56), and Cyt (57) facilitating multi-therapy resistance.

Notably, many candidates were involved in the metabolic rewiring/reprogramming mechanism that is a hallmark of cancer, allowing the cancer cells to alter their energy metabolism and associated pathways for survival (58). These included candidates such as PFKFB3, shown to be resistant for 10 out 20 drugs (59–66). Also, PKM (Pyruvate kinase M1/2) was resistant to 10 drugs including Dox (67), Cis (68), and so on (69–75). However, this scoring has limitations which are addressed in the discussion section.

### Cross-validation of molecules and mutations in our dataset and the available databases

To expand our analysis and identify the common drug-resistance markers, we compared our curated dataset with the molecules from existing drug-resistant databases such as DRESIS and DRMref. By cross-referencing our data with these databases, we found several overlapping molecules responsible for breast cancer drug resistance. We identified 385 molecules that were involved in breast cancer drug resistance from DRESIS db. Out of this, 140 were found common between this db and our dataset (**Supplementary** Figure 4a) including major candidates such as ABC transporters (ABCB1, ABCC2, etc), miRNAs (MIR125A, MIR340, etc), activators of survival pathways (STAT3, BIRC5, HIF1A, EGFR, etc) associated with 12 common anticancer drugs including Cap, FU, Tam, Dox, Vin, Ima, Eto, Pac, Cyc, Doc, Cis, and Tra.

From the DREMref db, there were 5086 molecules that were associated with breast cancer drug resistance from different datasets including BC cell lines, PBMC, and tumor tissues. 232 candidates were common between DREMref and our dataset (**Supplementary** Figure 4b) with 16 common drugs such as Rit, Bor, Gef, Gem, Cis, Vin, FU, Met, Ima, Tam, Eto, Dox, Pac, Dex, Doc, and Tra. Some of the overlapping candidates were BCL2, EGFR, TOP2A, AR, TUBB, ANXA1, among others. We also looked into the mutated genes from the COSMIC db. Around 428 genes were found to overlap between the BC patient and cell line COSMIC data and our dataset (**Supplementary** Figure 4c). 103 TCGA patients were also found to be common out of 106 from both the datasets (**Supplementary Table 11**).

### Development of the database and evaluation of its effectiveness in predicting drug resistance

In this study, we combined the information curated for the altered mRNAs, proteins, miRNAs, and lncRNAs showing drug resistance. This data was used to create a user-friendly Breast Cancer-Predict drug-resistant database (BC-Predict Database) that can be accessed through https://ciods.in/bcdrdb (**Figure 8**). The database is accessible through simple module including Drug Resistance Profiling. Based on the published cell line based studies, this module predicts the drug resistance pattern of the altered molecules given as a query. For this module, the users can provide the query based on the expression patterns such as downregulation or upregulation of a candidate (e.g. protein, miRNA). The outcome of this feature highlights the molecules within the query that have the potential to develop anticancer drug resistance. This also gives us information about multi-drug resistance associated with the altered molecule. Figure **9** demonstrates the effectiveness of this database by analyzing the expression profile as input for 1 poor prognosis (**Figure 9a**) and 1 good prognosis patients (**Figure 9b**). Here the differentially expressed proteins in post-treatment residual tumor vs pre-treatment tumor procured from the proteomics dataset of breast cancer patients, were used as input (76). Based on this, we could investigate the anticancer drug profiles to which the patients are more or less prone/probable to develop resistance. The information provided in this study for developing the database has to be further validated as this provides the groundwork for predictive analysis for translational purposes.

**Figure 8:**
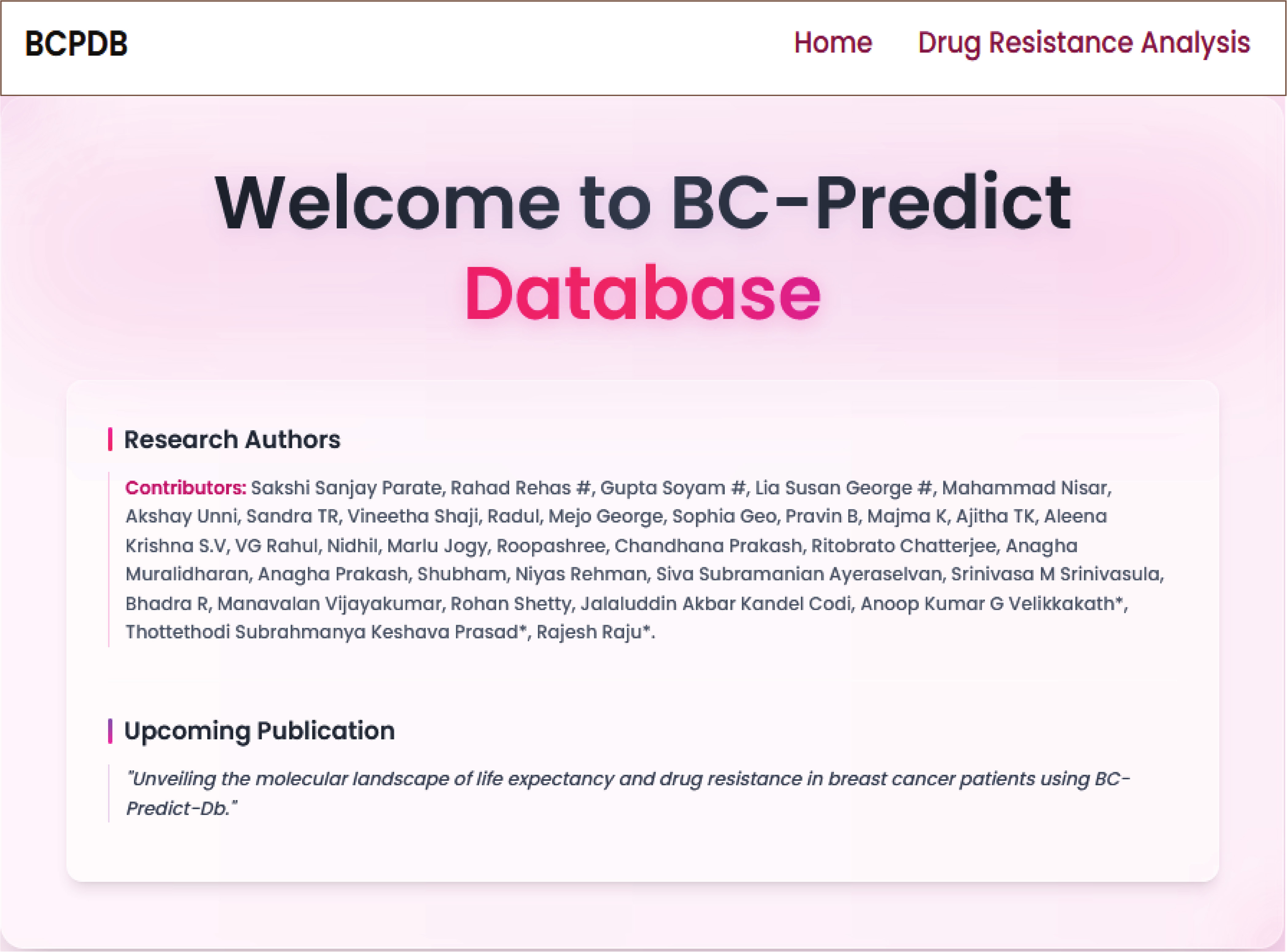
**Snapshot of the breast cancer drug resistance database (BC-Predict Database)**

**Figure 9.**
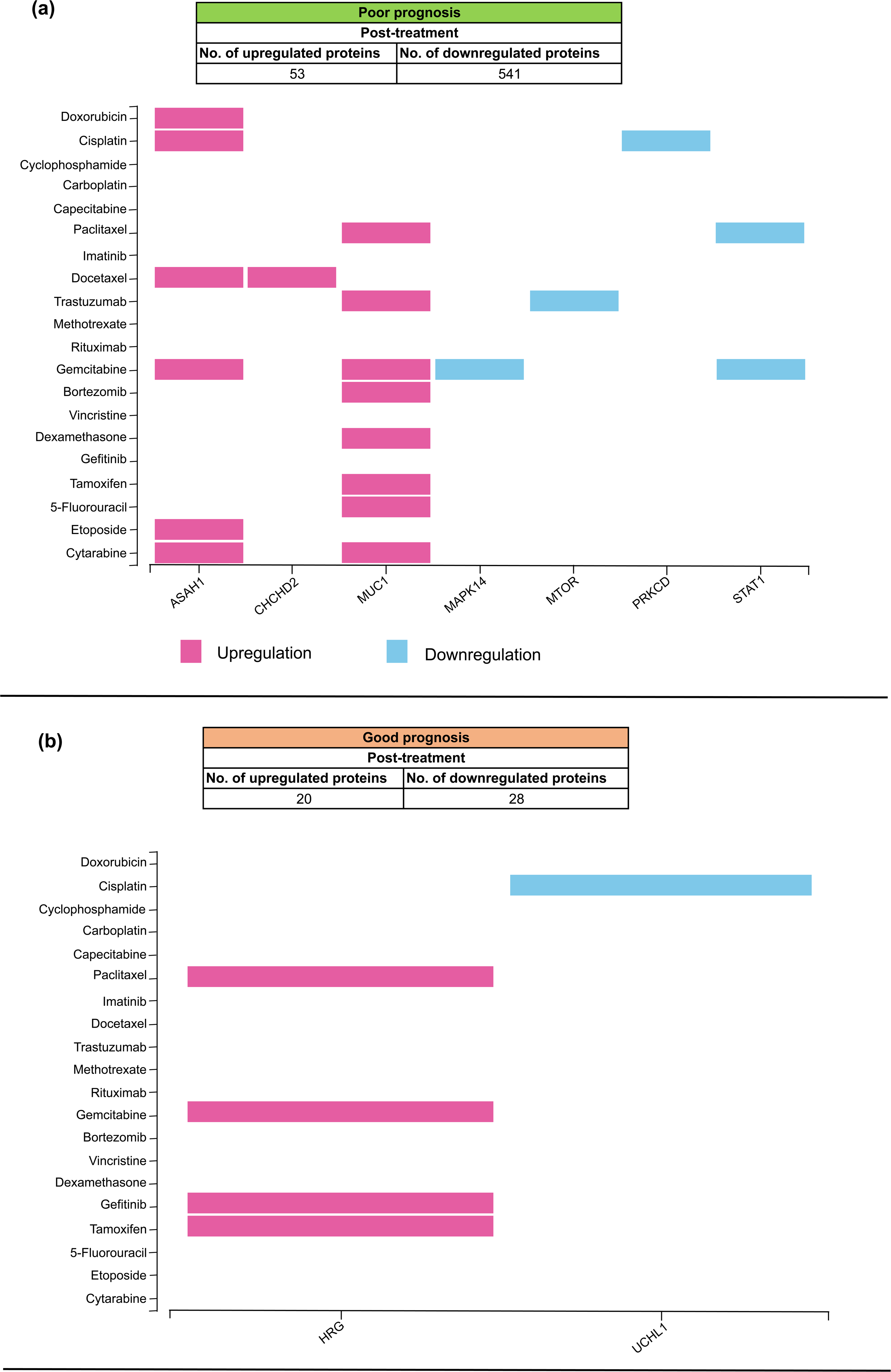
Snapshot of the drug resistance profile of differentially expressed proteins for validation of BC-Predict Database from the proteomics dataset of pre-treatment core needle biopsy vs post-treatment residual tumor samples of breast cancer patients. (**a**) Patient with poor prognosis; (**b**) Patient with good prognosis.

## Discussion

In our study we generated data for (1) mRNA candidates that distinguish deceased and survival groups of breast cancer patients, (2) Inventory of candidate mRNAs overexpressed in at least 80% breast cancer patients, (3) Inventory of breast cancer drug resistance related candidates, manually curated against 20 chemotherapy drugs (4) A database that could predict drug resistance and survival probability, (5) List of top priority drug resistance associated candidates that are involved in multidrug resistance (>15 drugs, 5-15 drugs, 2-4 drugs), (6) List of candidates that are involved in multidrug resistance in other cancers, however their drug resistance role for more than 15 chemotherapy drugs in the market are not studied and (7) their role in breast cancer drug resistance is not studied.

By analyzing 106 breast cancer patients from the cProsite, we identified a number of differentially expressed mRNAs in at least 80% of the patients. This analysis revealed that the upregulated mRNAs were predominantly involved in cell cycle regulation, cytoskeletal reorganization, DNA repair, and other cellular processes. Conversely, the downregulated mRNAs were majorly related to kinase signaling. These findings suggest that the involvement of some of these altered mRNAs may contribute to the development of drug resistance. Altered cell cycle regulation and defects in cell cycle checkpoints may lead to rapid proliferation and reduce the efficacy of target drugs, also allowing the damaged DNA to enter mitosis and further lead to genetic instability (77,78). These mRNAs may have the potential to be the drivers in drug resistance as they may be involved in tumor progression by activating oncogenic signaling pathways (79). The upregulated mRNAs in the deceased group that may be involved in drug resistance can be one of the reasons for their poor survival (80). This study also provides insights through understanding the mRNAs that distinguish deceased from surviving groups of breast cancer patients.

Apart from these, other molecules involved in drug resistance were also curated from the existing literature. By focusing on specific cell lines representing three breast cancer subtypes, namely, luminal, TNBC, and HER2-positive, we narrowed down our search to relevant literature. We found a pattern of multi-drug resistance of various molecules backed up by the literature involving their specific role in the drug resistance by either activation, downregulation, upregulation, silencing, or knock-out studies. These molecules were found to commonly target multiple anticancer drugs such as Pac, Cis, Dox, and so on, suggesting their pivotal role in drug resistance. A clear pattern was observed with some candidates frequently upregulated such as anti-apoptotic regulators (eg. BCL2 family, BIRC7, NFKB1, IL6), cell cycle regulators (eg. CDK6, CDKN1B, MDM2, CHEK1), drug efflux proteins (eg. ABC transporters), cell survival (eg. AKT1, MTOR, STAT3), and different signaling mechanisms including JAK-STAT, VEGF, MAPK, mTOR and so on. There were certain candidates showing consistent downregulation across different drugs including miRNAs regulating cancer pathways (eg. MIR130A, MIR520B), pro-apoptotic regulators (eg. APAF-1, BAX, BID), DNA repair genes (eg. RAD23A), and tumor suppressor genes (eg. PTEN), among few. Also, some candidates showed resistance pattern of upregulation to certain drugs while downregulation to other drugs depending on the cell type, drug action, and context, for eg. CASP3 when downregulated inhibits apoptosis (81) and when upregulated promotes apoptosis and resensitizes the cells to cell death (82), also play a role in promoting tumorigenesis by causing genomic instability (83). Our findings align with the existing research on the molecular drivers of drug resistance including the implication of drug efflux pumps such as ABC transporters (84), regulation of CSCs such as ABCB5, ALDH1, EGFR, enhancing EMT such as cytokeratins, cadherins, miRNAs (84,85), among other mechanisms.

In order to extend our understanding of the molecular mechanisms in drug resistance, we combined our findings with publicly available databases. By cross-referencing our dataset with DRESIS and DRMref databases, we identified common molecules associated with drug resistance across multiple studies. These include molecules such as RRM2 (86), BIRC5 (87), FGF2 (88), that are responsible for escaping apoptosis. Molecules such as EGFR (89) are involved in proliferation, RAD51 (90) involved in cancer progression, and RAD23A (91) involved in autophagy regulation. Some candidates such as CDC6 (92), AURKA (93), AURKB (94), MYBL2 (95), CDKN3 (96), CDK1 (97), TOP2A (98), NUSAP1 (99), among few, are responsible for cell cycle-mediated resistance to several combination treatments (77). These existing databases provide valuable resource by compiling large-scale studies including genomics and transcriptomics, also single-cell datasets that give a broad perspective of altered genes in drug resistant versus control cells. Consequently, the observation of simply upregulation and downregulation in the transcriptomic data does not provide definitive evidence of its contribution in conferring drug resistance (100). Therefore, in our curated dataset we narrowed our analysis to functional validation approaches including exclusive overexpression, silencing, knockout, or knockdown studies from published articles, that led us to confidently infer its role in anticancer drug resistance. Considering the clinical implications of this study by integrating the mRNA expression profiles with clinical outcomes, it offers a starting point to find out the likelihood of response to standard drugs and tailor their treatments accordingly (101,102). Using a multi-pronged approach by including feasible and affordable combination therapies, targeted therapies, and immunotherapies, may overcome anticancer drug resistance (103,104).

However, it is necessary to recognize the limitations of this study. If a particular protein has not been studied in the context of a specific drug, it was not scored in the matrix and not selected for prediction. Therefore, candidates showing no involvement in drug resistance (according to this database) might either indeed have no involvement in drug resistance, or they may have been a less explored candidate. Another limitation is that, if a protein is involved in drug resistance in other cancer cell lines, however, its role in breast cancer is not tested, we still considered it, as it cannot be ignored. This is because it is likely that a protein causing drug resistance in another cell line may also show drug resistance in breast cancer cell lines, even though there could be exceptions. The small sample size of 106 patients is not sufficient to fully capture the molecular heterogeneity of breast cancer. By checking the individual clinical data from TCGA, it was found that the treatment data for the deceased patients was not available. It may be that these patients were not given any therapy or the treatment data was not recorded or concealed. Furthermore, the mRNA expression data from publicly available datasets, while valuable, may not fully reflect the complexities of *in vivo* tumor biology (105,106),(107). There were various molecules for which drug resistance studies were not available. Although the data biocuration was focused on breast cancer, we also included studies with different cancer types. The drug resistance pathways may share some similarities in various malignancies (108), but it is crucial to acknowledge that the specific context of breast cancer may affect the relevance and applicability of these findings.

To further confirm the functional significance of the molecules and their role in breast cancer drug resistance, more validation studies are needed. The drugs were chosen for biocuration based on their clinical relevance and use in breast cancer treatment. This analysis focused on the generic drug names for consistency; however, the particular brand names of the same drug were not taken into account. Further investigations can improve these by considering the impact of specific drug brands and formulations. From our analysis, it is revealed that the large-scale protein overexpression studies (focused on candidates involved in drug resistance) done in breast cancer cell lines focusing on drug resistance do not reflect the overexpressed mRNA profiles of real life patients. This claim is made based on the observation that the mRNAs overexpressed in breast cancer patients did not overlap much with the candidates studied for drug resistance in cell line based studies in publications. However, it should also be noted that mRNA expression profiles may not always necessarily match with protein expression profiles. Future studies focusing on drug resistance predictions shall focus on filling the gaps mentioned above.

The molecular markers identified here may contribute to the development of personalized treatments especially when gene editing based therapeutics are making it to the forefront successfully.

## Availability of data and materials

The cProsite website (https://cprosite.ccr.cancer.gov/), TCGA website (https://www.cancer.gov/tcga) for patient data. DRESIS db (https://dresis.idrblab.net/), DRMref db (https://ccsm.uth.edu/DRMref/), COSMIC db (https://cancer.sanger.ac.uk/cosmic). BC-Predict-Db (https://ciods.in/bcdrdb), Proteomic dataset (PMID: 32960509; (project number: PXD012000; http://www.ebi.ac.uk/pride/archive/projects/PXD012000).

## CRediT authorship contribution statement

**SSP:** Methodology, Data curation, Formal analysis, Visualization, Writing- Original Draft, Writing- Review & Editing. **RR:** Methodology, Software, Data curation, Writing- Original Draft, Cross-validation, Writing- Review & Editing. **GS:** Methodology, Data curation, Writing- Original Draft, Writing- Review & Editing. **LSG:** Methodology, Data curation, Writing- Original Draft, Writing- Review & Editing. **MN:** Methodology, Software, Data curation, Validation,

Writing- Original Draft, Writing- Review & Editing. **AU:** Methodology, Data curation, Writing- Original Draft, Writing- Review & Editing, Validation. **STR:** Methodology, Data curation, Validation, Writing- Original Draft, Writing- Review & Editing. **SM:** Data curation, Validation, Formal analysis, Visualization, Writing- Original Draft & Editing. **VS:** Methodology, Software, Formal analysis, Visualization, Validation. **AKSV:** Methodology, Data curation, Writing- Original Draft, Writing- Review & Editing, Validation. **MG:** Methodology, Visualization, Writing- Original Draft, Writing- Review & Editing. **BR:** Methodology, Data curation, Writing- Original Draft, Writing- Review & Editing, Validation. **RRD:** Methodology, Software, Writing- Original Draft, Writing- Review & Editing. **PB:** Methodology, Software, Data curation, Validation, Writing- Original Draft, Writing- & Editing. **SSA:** Methodology, Software, Data curation, Writing- Original Draft, Writing- Review & Editing. **MK:** Methodology, Data curation, Validation, Writing- Original Draft, Writing- Review & Editing. **ATK:** Methodology, Formal analysis, Writing- Original Draft, Writing- Review & Editing. **RC:** Methodology, Data curation, Validation, Writing- Original Draft, Writing- Review & Editing. **RVG:** Methodology, Data curation, Validation, Writing- Original Draft, Writing- Review & Editing. **MJ:** Methodology, Data curation, Validation, Writing- Original Draft, Writing- Review & Editing. **RPG:** Methodology, Data curation, Validation, Writing- Original Draft, Writing- Review & Editing. **CP:** Methodology, Data curation, Writing- Original Draft, Writing- Review & Editing. **AMu:** Methodology, Data curation, Writing- Original Draft, Writing- Review & Editing. **APr:** Methodology, Data curation, Writing- Original Draft, Writing- Review & Editing. **SSU:** Methodology, Formal analysis, Writing- Original Draft, Writing- Review & Editing. **AA:** Methodology, Data curation, Writing- Original Draft, Writing- Review & Editing. **NR:** Methodology, Software, Writing- Original Draft, Writing- Review & Editing. **MV:** Supervision, Resources, Writing- Original Draft, Writing- Review & Editing. **RS:** Supervision, Resources, Writing- Original Draft, Writing- Review & Editing. **JAKC:** Supervision, Resources, Writing- Original Draft, Writing- Review & Editing. **AKGV:** Conceptualization, Supervision, Data curation, Validation, Formal analysis, Visualization, Writing- Original draft, Validation, Writing- Review & Editing. **TSKP:** Conceptualization, Methodology, Resources, Writing - Review & Editing, Supervision, Resources. **RRj:** Conceptualization, Methodology, Supervision, Resources, Writing- Review & Editing. All the authors participated in drafting and reviewing the manuscript. All the authors read and approved all contents of the final manuscript.

## Corresponding authors

Thottethodi Subrahmanya Keshava Prasad, Anoop Kumar G. Velikkakath, Rajesh Raju.

## Acknowledgments

We thank Yenepoya (Deemed to be University) for its support in establishing the Centre for Integrative Omics Data Science (CIODS). The authors acknowledge the support of the Department of Biotechnology, Government of India to the Yenepoya (Deemed to be University) through the project on “Skill Development in Mass Spectrometry-based metabolomics technology BIC” (BT/PR40202/BTIS/137/53/2023). We thank Karnataka Biotechnology and Information Technology Services (KBITS), the Government of Karnataka, for the infrastructure support of the Center for Systems Biology and Molecular Medicine at Yenepoya (Deemed to be University) under the Biotechnology Skill Enhancement Program in Multiomics Technology (BiSEP GO ITD 02 MDA 2017).

## Funding

This research did not receive any specific grant from funding agencies in the public, commercial, or not-for-profit sectors.

## Ethical declarations

Not applicable

## Conflict of interests

The authors declare that they have no conflict of interest.

## Additional information

**Disclaimer:** The authors acknowledge any discrepancies in the data biocuration. We encourage the readers to contact the corresponding authors regarding any concerns that may arise. This will allow us to promptly investigate and address any identified issues in a timely and transparent manner and ensure that data integrity is maintained.

## Notes

### Competing Interest Statement

The authors have declared no competing interest.

