## Supplementary Figures 1-4 for "BC-Predict Database: A Curated Resource of Experimentally Validated Markers in Multidrug Resistance in Breast Cancer"

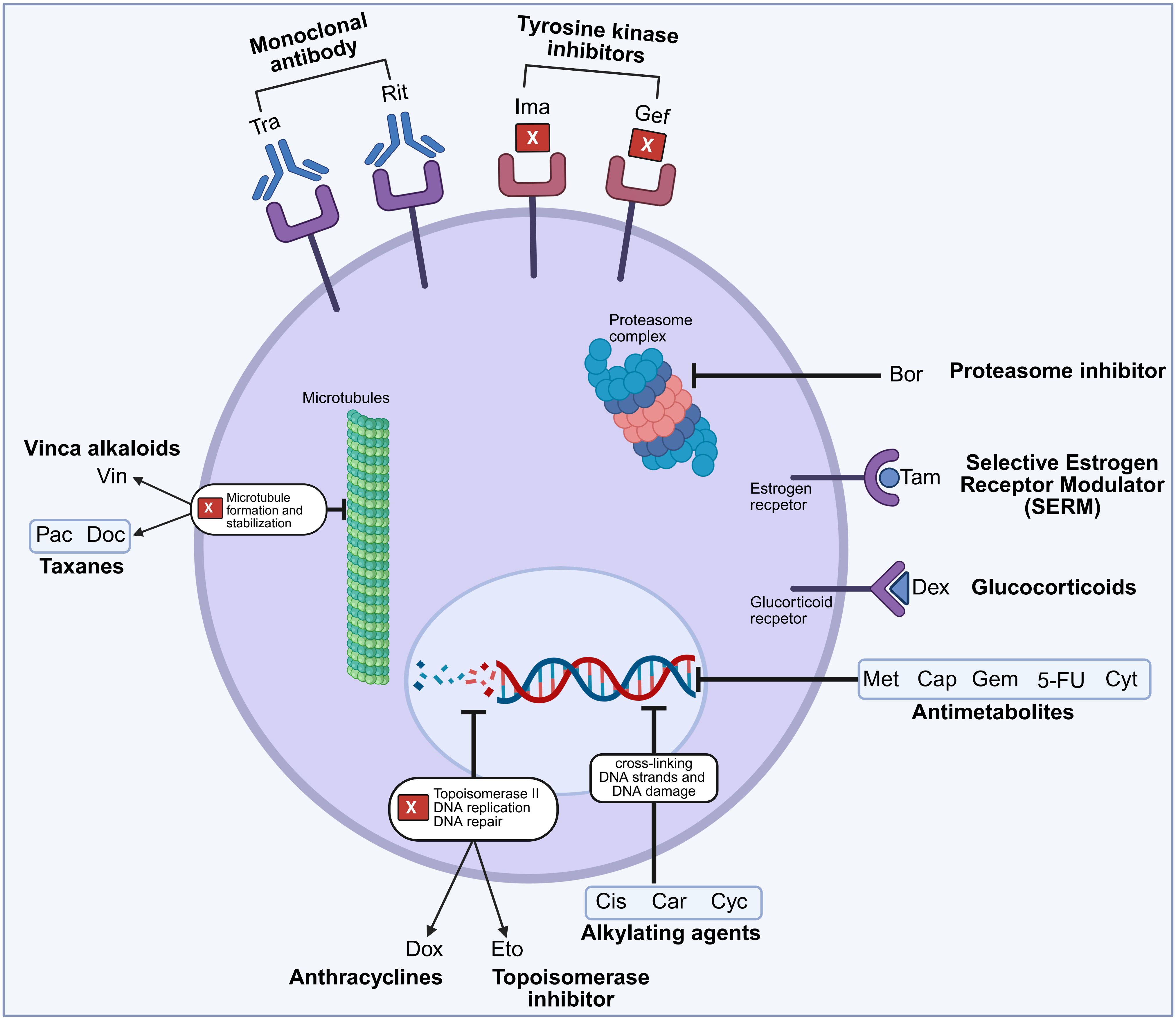


**Abbr:**
Tra- Trastuzumab
Rit: Rituximab
Ima: Imatinib
Gefitinib
Bor: Bortezomib
Tam: Tamoxifen
Dex: Dexamethasone

Met: Methotrexate
Cap: Capecitabine
Gem: Gemcitabine
5-FU: 5-Fluorouracil
Cyt: Cytarabine
Cis: Cisplatin

Car: Carboplatin
Cyc: Cyclophosphamide
Eto: Etoposide
Dox: Doxorubicin
Pac: Paclitaxel
Doc: Docetaxel
Vin: Vincristine

**Supplementary Figure 1.** An overall representation of different classes of anticancer drugs, their targets, and cytotoxic effects on the cancer cell.

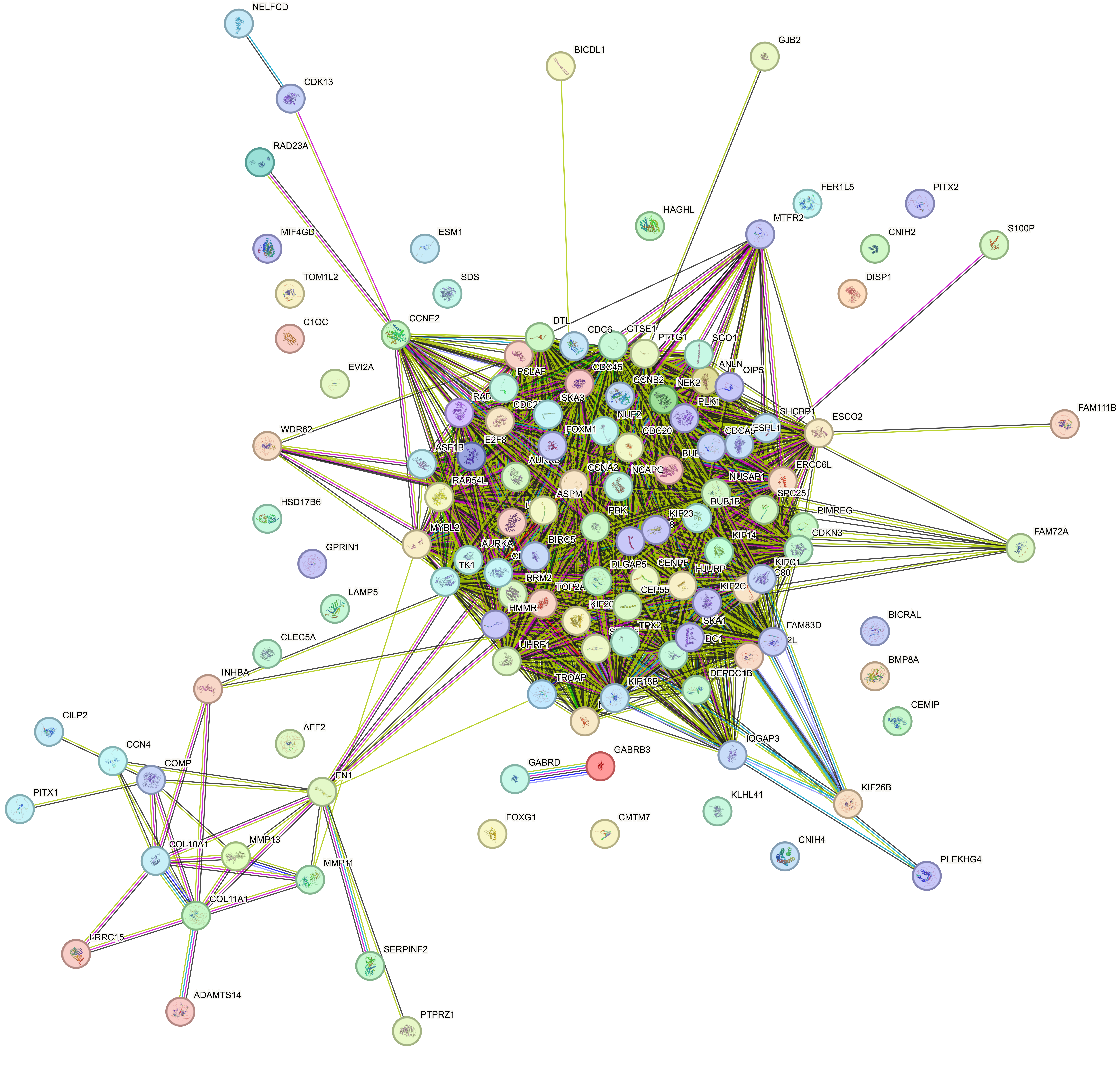


**Supplementary Figure 2:** Network analysis using STRING db of the commonly overexpressed mRNAs in at least 80% of breast cancer patients

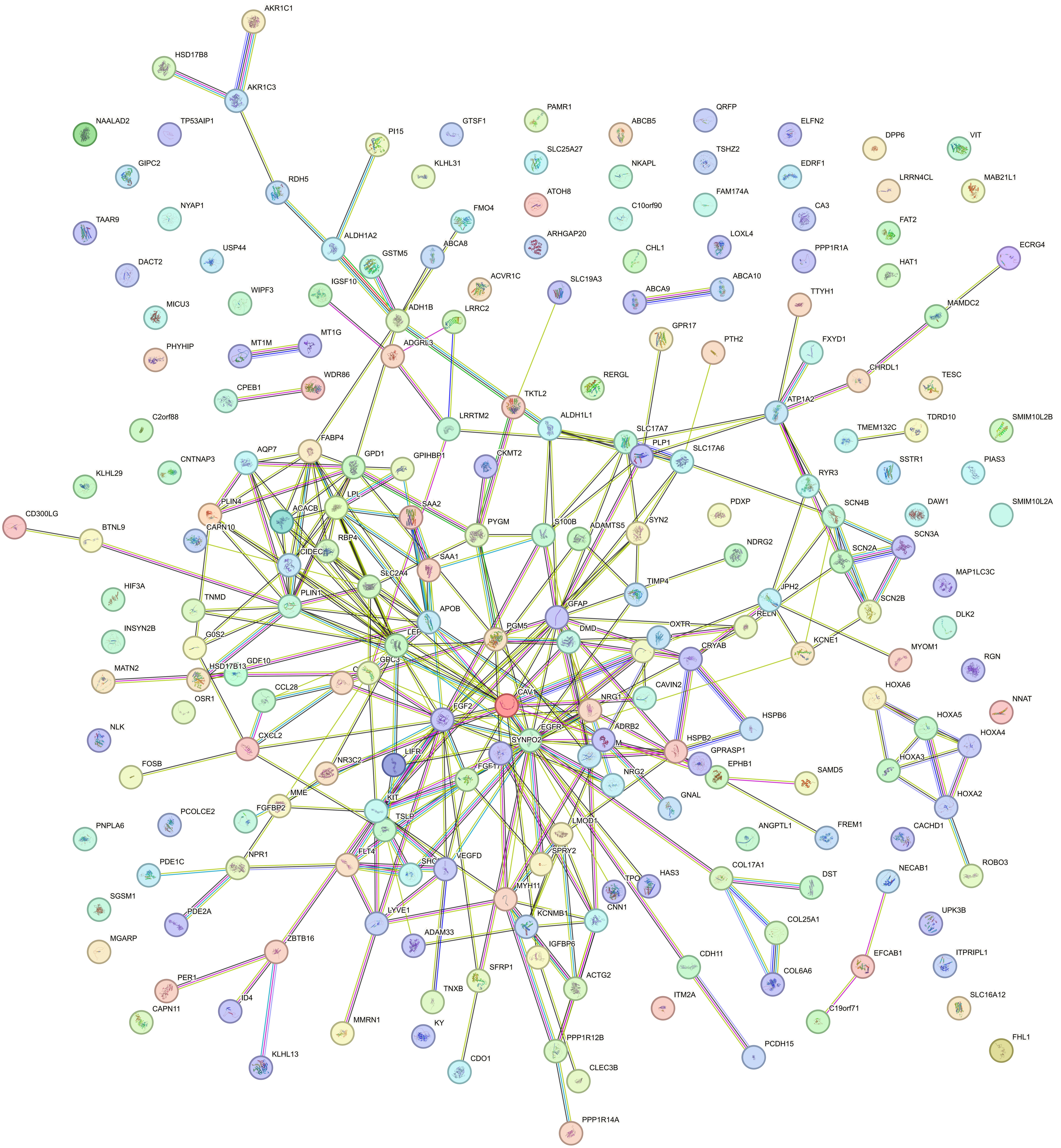


**Supplementary Figure 3:** Network analysis using STRING db of the commonly downregulated mRNAs in at least 80% of breast cancer patients

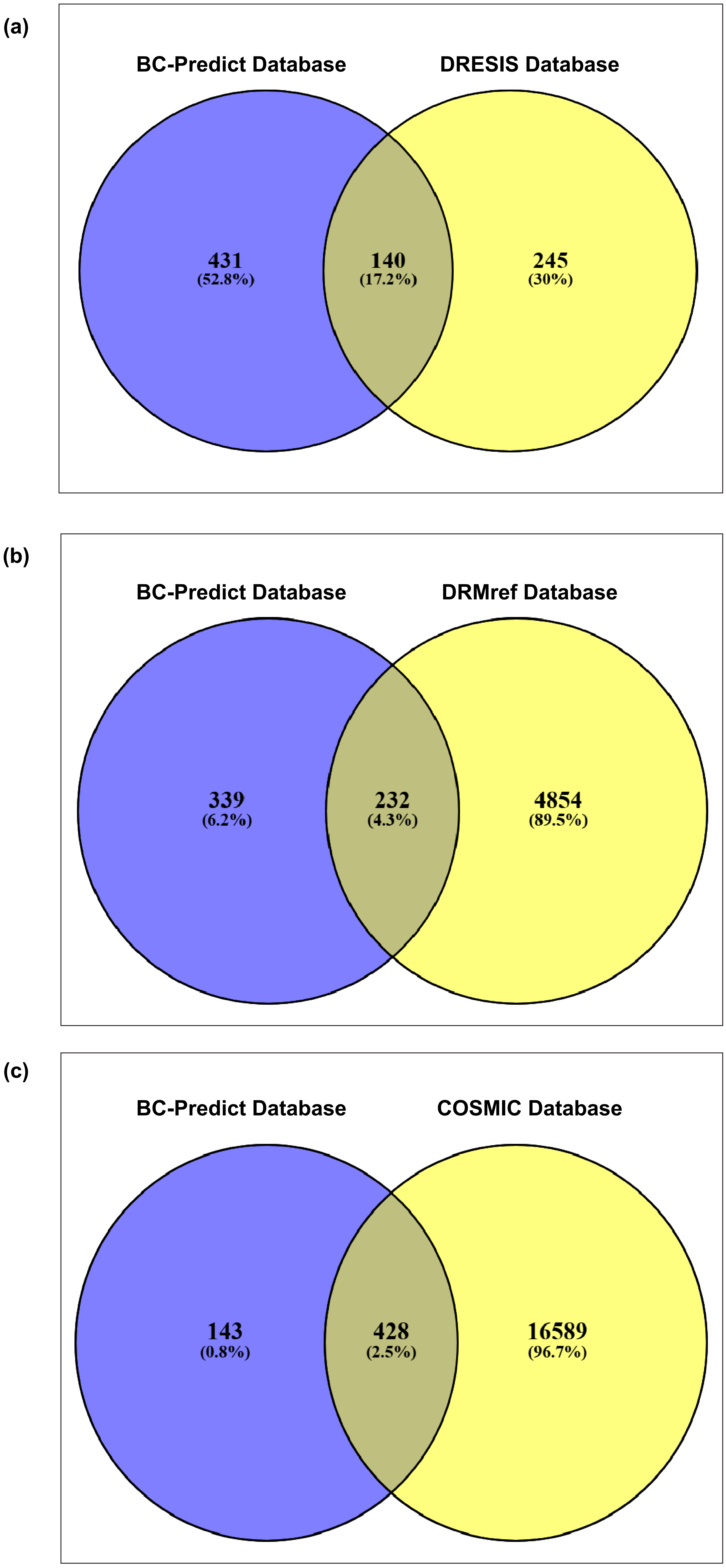


**Supplementary Figure 4.** Venny result of the comparison of molecules from the existing drug resistance databases and our curated dataset. (a) DRESIS database versus BC-Predict Database, (b) DRMref database versus BC-Predict Database, and (c) COSMIC database versus BC-Predict Database
